# Language-Informed Basecalling Architecture for Nanopore Direct RNA Sequencing

**DOI:** 10.1101/2022.10.19.512968

**Authors:** Alexandra Sneddon, Nikolay Shirokikh, Eduardo Eyras

**Affiliations:** Australian National University

## Abstract

Algorithms developed for basecalling Nanopore signals have primarily focused on DNA to date and utilise the raw signal as the only input. However, it is known that messenger RNA (mRNA), which dominates Nanopore direct RNA (dRNA) sequencing libraries, contains specific nucleotide patterns that are implicitly encoded in the Nanopore signals since RNA is always sequenced from the 3’ to 5’ direction. In this study we present an approach to exploit the sequence biases in mRNA as an additional input to dRNA basecalling. We developed a probabilistic model of mRNA language and propose a modified CTC beam search decoding algorithm to conditionally incorporate the language model during basecalling. Our findings demonstrate that inclusion of mRNA language is able to guide CTC beam search decoding towards the more probable nucleotide sequence. We also propose a time efficient approach to decoding variable length nanopore signals. This work provides the first demonstration of the potential for biological language to inform Nanopore basecalling. Code is available at: https://github.com/comprna/radian.

## 1 Introduction

A crucial mechanism underpinning gene expression in eukaryotes is alternative splicing. It is a process in which exons in precursor RNA molecules are differentially included in mature messenger RNA (mRNA) transcripts, which endows each gene with the potential to give rise to an array of transcript isoforms[1]. Splicing is known to be intrinsically linked to both normal and pathological cellular function, thus a means to delineate the subtle variations between splicing patterns in different biological contexts is essential[2, 3]. An obvious yet challenging pre-requisite is the ability to sequence mRNA transcripts with high accuracy.

The sequencing of full-length RNA molecules directly is possible with Oxford Nanopore Technologies’ long-read direct RNA (dRNA) sequencing platforms. The technology utilizes a biological nanopore, through which an RNA strand translocates, producing an electrical signal that can be decoded into the nucleotide sequence (’basecalling’)[4, 5]. Basecalling is a challenging computational problem. At any given time, multiple nucleotides (a ‘k-mer’) occupy the nanopore, which means that each sample is influenced by *k* bases. For pore version R9.4, *k* is approximately 5, thus there are 4^5^ = 1024 possible k-mers that influence the current[5]. Each k-mer gives rise to a distribution of current levels and the distributions for different k-mers overlap[6]. In addition, the speed of translocation of the molecule through the pore is nonuniform[6] and noise associated with the nanopore sensing system further obscures the signal[7]. As a result, varied signals arise from sequencing the same molecule[6].

Basecalling accuracy for dRNA lags behind that for DNA, where basecalling developments to date have predominantly been focused[8–14]. Broadly, the field has moved towards convolutional neural networks[13–15] in place of recurrent architectures[8–10]. The practical advantage of this is improved efficiency with minimal effort, since the convolution operation can be easily parallelized in contrast to the sequential processing performed by recurrent networks. Importantly, stacked convolutional layers are able to capture the variance in the number of samples per nucleotide[16]. To incorporate information in the signal from neighbouring nucleotides, approaches have included the use of dilated convolutions[13], increasing the kernel size with network depth[16] and the inclusion of self-attention layers[14]. Recent work has recognised that an RNA-specific architecture is warranted given the differences between dRNA and DNA signals, namely the slow translocation speed (70 bases per second (bps) for dRNA in contrast with 450 bps for DNA)[16]. However, the nucleotide error rate remains at approximately 7% for the human transcriptome[16].

Not only do the signals differ, but from the perspective of nanopore sequencing the mRNA sequences themselves contain biases that are not present in DNA sequences. mRNA is known to contain specific nucleotide patterns[17] and since dRNA sequencing always takes place from the 3’ to 5’ direction of the molecule, the content of mRNA reads is ordered as follows: (1) poly-A tail, which is comprised of an adenine homopolymer, (2) 3’ untranslated region (UTR), (3) open reading frame (ORF), (4) 5’ UTR, (5) 7-methylguanosine cap.[18] Patterns can also be observed within each region of mRNA. For instance, the ORF is framed by a start codon (typically AUG) and stop codon (UAG, UAA or UGA), encoding the amino acid sequence of a protein with a trinucleotide periodicity.[18, 19] As such, dRNA reads repeatedly contain a specific sequence structure that is expected to be identifiable from the signal. In contrast, DNA may be sequenced in either the 5’ to 3’ or 3’ to 5’ direction at any point in the genome, thus the same pattern is not observed in nanopore DNA reads.

Since the majority of RNA in a dRNA sequencing library is mRNA (>80%), we hypothesized that it is possible to leverage the unique content and patterns in mRNA sequences as an additional layer of information during dRNA basecalling. In this work, we propose an approach to incorporate an mRNA language model into dRNA basecalling. **Our contributions are as follows: (1)** We devise a method to decode variable length nanopore basecalling signals, which we show is more efficient than the standard approach both empirically and through time complexity analysis. **(2)** We developed a probabilistic model of mRNA language that predicts the next nucleotide given a preceding sequence. **(3)** We modified the CTC beam search decoding algorithm to conditionally incorporate an mRNA language model. **(4)** We show that the mRNA language model is able to guide CTC beam search decoding towards the more probable nucleotide to provide an overall modest improvement to basecalling accuracy. This work highlights the potential for the inclusion of RNA language information to improve dRNA basecalling and provides a framework upon which future language-informed basecalling studies can build.

## 2 Methods

The high-level workflow for our language-informed basecalling architecture is depicted in **Fig. 1**.

**Figure 1:**
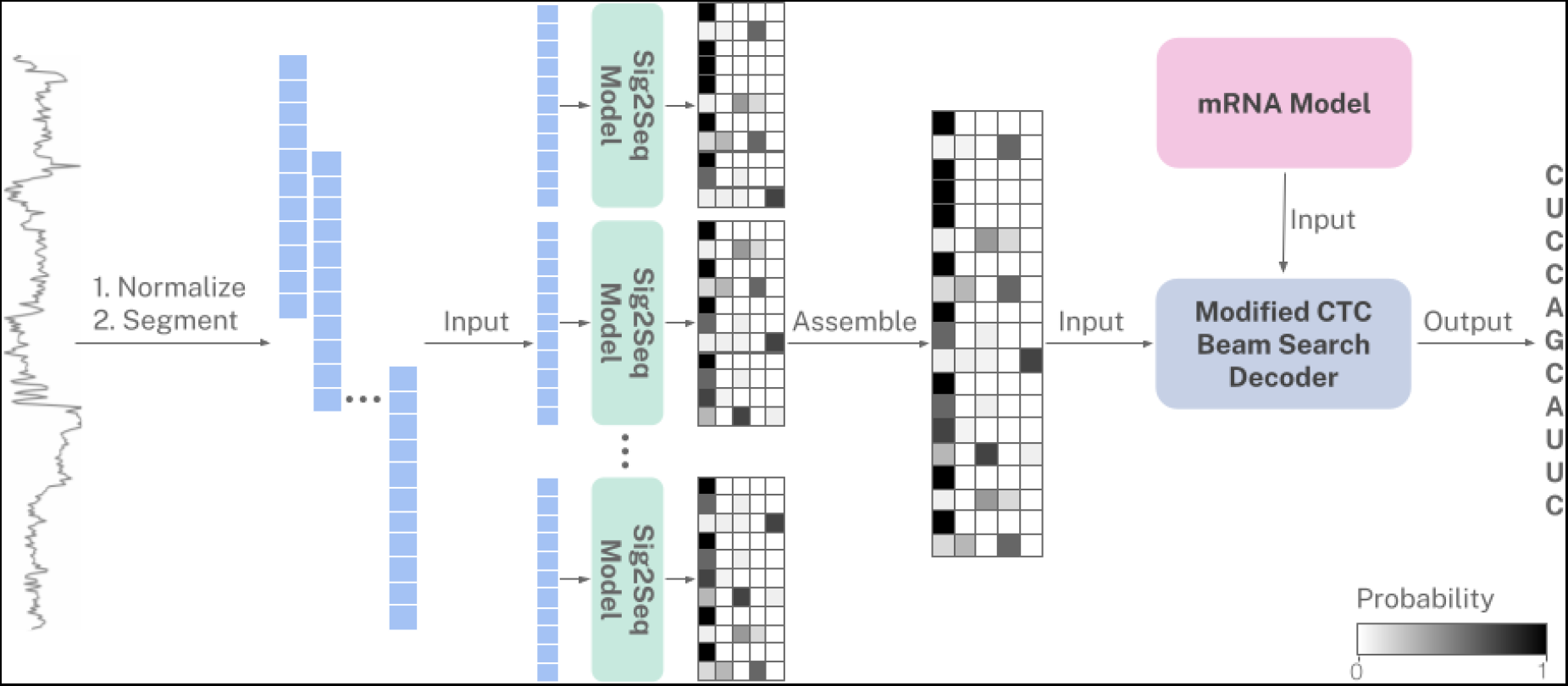
End-to-end basecalling architecture. Raw nanopore signals are normalised using median absolute deviation with outlier smoothing, then segmented into overlapping chunks of length 1024 with step size 128. Each chunk is fed to the signal to sequence (Sig2Seq) model, which outputs a CTC matrix comprising a probability distribution across nucleotides for each timestep. The ‘chunk-level’ CTC matrices are assembled to produce a ‘global’ CTC matrix covering the entire input signal. The nucleotide sequence is then elucidated using CTC beam search decoding, modified to allow dynamic inclusion of an mRNA model, which encodes the likelihood of each nucleotide given a preceding nucleotide sequence.

### 2.1 Signal to Sequence Model

#### 2.1.1 Training Data

Raw nanopore signals from two human heart and HEK293 MinION Mk1B dRNA sequencing runs were used to train the signal to sequence (Sig2Seq) model. Heart fast5 files were obtained from the European Nucleotide Archive under the accession PRJEB40410[20], while HEK293 fast5 files were generated as part of this study (see **??**). The approach used to generate labelled signal chunks for training the Sig2Seq model is detailed in **??**. 20% of the chunks in each training set were reserved for validation.

#### 2.1.2 Model Development

Basecalling is an order-synchronous but time-asynchronous sequence to sequence classification problem, where the objective is to map a nanopore signal of length *N* to a nucleotide sequence of length *M*, where *M* ≤ *N*. We trained a deep neural network to learn the transformation from signal to sequence using connectionist temporal classification (CTC)[21], which treats the network output as a conditional probability distribution over all possible sequences. CTC loss was the objective function used during training, which was optimized using Adam[22] with an initial learning rate of 0.0001. Chunks were fed to the network in batches of size 32 and after each epoch the training set was shuffled and the network was evaluated using the validation set to avoid overfitting. Training was distributed across 4 NVIDIA Tesla V100 GPUs.

#### 2.1.3 Chosen Architecture

Since the objective of this study was to develop an algorithm for integrating language models during basecalling, for our Sig2Seq model we simply adapted a previously published DNA basecalling architecture, CausalCall. CausalCall[13] is based on a Temporal Convolutional Network (TCN)[23]. TCNs were specifically designed as an alternative to recurrent architectures for sequential data. They utilize dilated causal convolutions to allow the receptive field across past inputs to increase with network depth. Coupling dilated causal convolutions with residual connections allows very deep networks with a large “memory” to be trained.[23] TCNs have been shown to outperform equivalent-capacity recurrent architectures with greater efficiency on a range of standard sequence modelling benchmarks.[23]

Our Sig2Seq model architecture (**Fig. 2**) is based on CausalCall, but with a few key differences: (1) since RNA translocates slower than DNA, there are typically more samples per nucleotide so we increased the network depth and therefore the receptive field size, (2) we implemented the original residual block as previously described [23], (3) for training we used a much lower initial learning rate of 0.0001. Briefly, our model is comprised of six stacked residual blocks (kernel size *k* = 3, # filters *n* = 256, dilations *d* = [1, 2, 4, 8, 16, 32]) that extract local temporal dependencies from the signal. Two fully connected layers (# neurons *n* = 5, 128 respectively) then combine the features extracted from each channel. Finally, a softmax function converts the output for each timestep into a probability distribution across the possible nucleotides. The model was implemented using Keras[24] and has 2,200,581 parameters. The TCN layers were implemented using the keras-tcn Python library[25].

**Figure 2:**
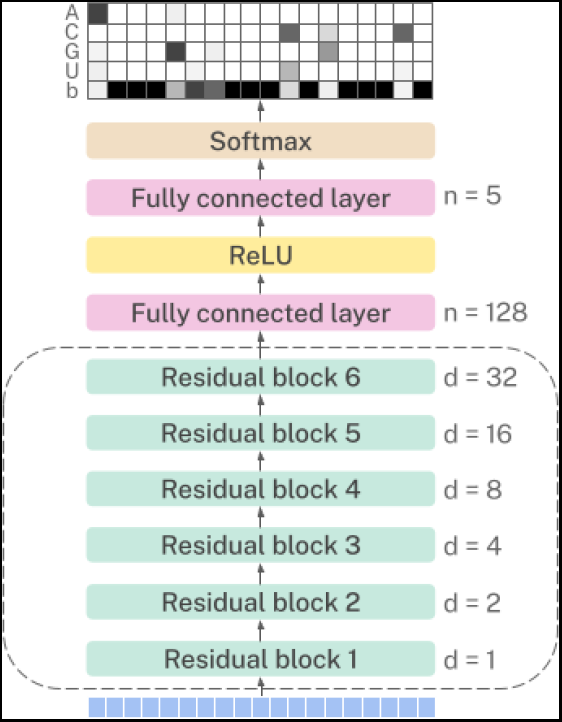
Architecture of the Sig2Seq model.

### 2.2 Language-Informed Decoding

#### 2.2.1 Assembly of Neural Network Outputs

For each input chunk, the Sig2Seq model outputs a 2D matrix with a row for each input timestep and column for each nucleotide, with an additional column for the blank symbol utilised by CTC. When the segmentation step size is less than the chunk length, each timestep in the input appears in multiple chunks (except for the timesteps at the start and end of the signal which appear only in the boundary chunks). To obtain the final basecalled sequence, existing basecallers either decode the CTC matrix corresponding to each chunk and then assemble the partial reads in sequence space[10, 13, 14, 16] or concatenate slices of the ‘chunk-level’ CTC matrices into a ‘global’ matrix before decoding, retaining the probability distribution from only one chunk-level matrix per timestep.[15]

The former capitalises on the network classifying each part of the signal multiple times in different contexts, since the chunks overlap. However, the expensive CTC decode operation is also performed per chunk and therefore multiple times for each part of the signal. In addition, since the order of partial reads is known, assembly of nucleotides is straightforward but may introduce errors due to the uncertainty of which nucleotides overlap between adjacent partial reads. In contrast, the latter approach precisely assembles the chunk-level matrices with efficient decoding by a single pass through the global matrix. However, since each row in the global matrix is sourced from only one chunk-level matrix, information from the remaining chunks at that position is lost.

We instead assemble the chunk-level CTC matrices by averaging the probability distributions for each timestep across chunks. Assembly is therefore precise but also retains information from all chunks per timestep. Importantly for this study, since CTC decoding is performed on the global matrix, the decoded path through the matrix corresponds to the nucleotide sequence from the beginning of the read. It therefore becomes possible during decoding to utilise knowledge of the sequence observed thus far.

#### 2.2.2 Modified CTC Beam Search Decoding

The final nucleotide sequence is obtained by decoding the CTC matrix. It is important to note that networks trained with CTC loss output null predictions (‘blanks’) between elements in the sequence to allow repeats to be encoded. For RNA basecalling the set of possible output symbols is therefore *S* = {*A, C, G, U, b*}, where *b* is the blank symbol. Given a sequence of CTC symbols, the final sequence is obtained by first collapsing adjacent repeated symbols and then removing blanks.[21] A naive approach to decoding a CTC matrix is to simply select the symbol with highest probability at each timestep. However, this is not guaranteed to find the most likely final sequence since it does not take into account that there may be multiple paths corresponding to the same sequence.[26] For instance, “A-ACA”, “AA-ACA”, “A–ACCA” all reduce to “AACA” after collapsing repeats and removing blanks, but a greedy approach will treat these paths separately.

Instead, a beam search algorithm was proposed in [26], which keeps track of the *w* best paths at each step of the decoding, where *w* is the beam width. At each step of the decoding, the paths are sorted, the *w* best paths are selected and each extended by each symbol in *S*. The algorithm also caters for inclusion of a language model for automatic speech recognition (ASR), where the objective is to transform an audio signal into text. In ASR, the language model is used for computing the probability of the text corresponding to each path, so each time the path is extended with a new symbol, the text score of the path is updated using the language model. When sorting the paths at each timestep, the text score and path scores are combined to give the ranking.

Here we propose an alternative approach for conditionally incorporating a language model directly with the CTC probabilities. In basecalling it is important to accurately call every nucleotide since variations even in single nucleotides can be biologically significant. It is therefore important for the basecalled sequences to derive as much information as possible from the nanopore signal. However, in cases where the signal noise cannot be confidently resolved, we hypothesize that a language model will be of benefit in guiding classification towards the most likely nucleotide. Our proposed approach involves incorporating the mRNA model at each beam extend step. As a measure of how confident the prediction directly from the signal is at a given timestep *t*, we compute the entropy of the distribution across *S* predicted by the Sig2Seq model at *t*. If the entropy of the distribution exceeds a threshold *e^s^*, the signal is deemed sufficiently unresolved and a candidate for including the mRNA model. The mRNA model then predicts the probability across the four possible nucleotides *S* − *b* given the preceding *k* nucleotides in the current beam. We employ another entropy threshold *e^r^* and if the entropy of the mRNA model prediction is less than *e^r^* we average the Sig2Seq and mRNA model predictions across *S* − *b*.

**Algorithm 1** lists the pseudocode for our modified CTC beam search, which is based on [26]. Highlighted red and green are the steps we have removed and added, respectively. The inputs to the algorithm are CTC matrix *m*, beam width *w*, RNA model *r*, context length *k* and signal and mRNA model entropy thresholds *e^s^* and *e^r^*, respectively. *P_b_* and *P_nb_* are the probabilities a beam ends in a blank or non-blank, respectively.

### 2.3 mRNA Model

The objective of the mRNA model is to estimate the probability of the next nucleotide in the basecalled sequence, given the nucleotides already called. Since the ORF of mRNA encodes for an amino acid sequence, we expect strong trinucleotide biases in the coding sequence as well as regulatory sequence biases localised to the flanking UTRs[19], thus we expect short-range patterns in the mRNA sequence to have strong predictive power. We therefore assumed the Markov property that the probability of a given nucleotide depends only on a small number of immediately preceding nucleotides and developed a probabilistic model based on k-mers. We define a sequence of *k* nucleotides (a k-mer) as *n*_1:*k*_ and the joint probability of each nucleotide in the k-mer having a certain value as *P* (*n*_1:*k*_). The k-mer model estimates the probability of a nucleotide *n_i_* conditioned on the preceding *k* − 1 nucleotides as:

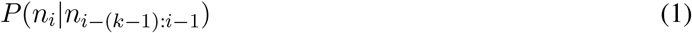

**Algorithm 1.**
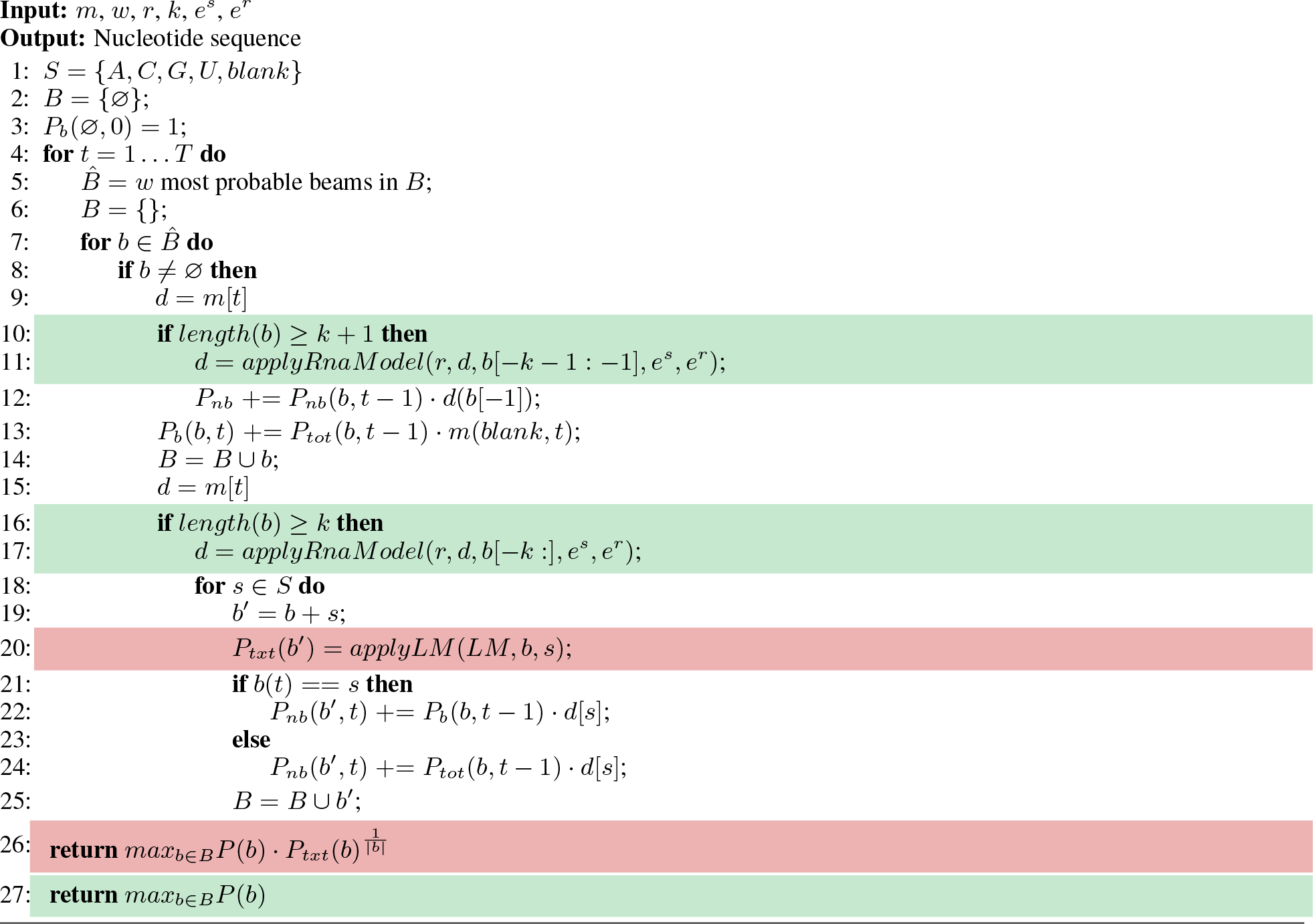
Modified CTC Beam Search

Maximum Likelihood Estimation was used to estimate the parameters of the k-mer model that maximise the likelihood of the k-mer frequencies in the protein-coding transcriptome given the model. Starting with all protein-coding transcripts in the human reference transcriptome (GRCh38.p13 release 34), excluding duplicate sequences, we reversed all sequences to a 3’ to 5’ orientation to match the dRNA sequencing direction and then counted k-mers using Jellyfish *count*[27] with options *−s 380M −t 10* in conjunction with either *-m 6* or *-m 12* to count 6-mers or 12-mers, respectively. We chose two values for *k* to allow comparison between different context lengths, with multiples of 3 selected since we expect trinucleotide biases in the ORF. 12 was the maximum value of *k* with a reasonable compute time. We estimated the conditional probability of a nucleotide *n_i_* given the preceding *k −* 1 nucleotides by normalizing the count *C* of the k-mer *n*_1:*k*_ by the count of all k-mers starting with the (k-1)-mer *n*_1:*k*−1_ as follows:

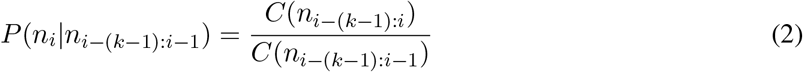

The k-mer models were evaluated extrinsically on the downstream basecalling task.

### 2.4 Evaluation

After optimization of the decoding hyperparameters (see **??**) basecalling performance was evaluated using reads from the nanopore Whole Genome Sequencing Consortium GM12878 project[28], Johns Hopkins University Run 1 (kit SQK-RNA001, pore R9.4). The reads were basecalled, mapped to the reference transcriptome and alignments filtered using the same approach described in **2.1.1**. 5,000 reads that aligned to protein-coding transcripts were then randomly selected to use for testing. The test reads were basecalled end-to-end following the workflow depicted in **Fig. 1** using the optimal set of decoding hyperparameters (*k* = 12, *e_s_* = 0.5, *e_m_* = 0.5) and a beam width of 6. To evaluate the effect of the RNA model, the test reads were basecalled again but the RNA model was excluded from decoding.

Following the standard basecaller evaluation approach[13, 14, 16], the basecalled reads were aligned to the transcriptome (GRCh38.p13 release 34) by minimap2[29] (v2.17) with options *−ax map-ont −uf –secondary=no −t 15*. The alignment CIGAR strings were parsed and basecalling accuracy evaluated using the sequence identity rate 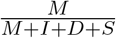 and total error rate 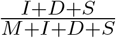, where *M*, *I*, *D* and *S* are the number of matches, insertions, deletions and substitutions, respectively. Since we filtered the test reads to only include those with high-confidence assignments to protein-coding transcripts, we knew the portions of the reference transcripts that the test reads align to. This enabled us to also align our basecalled reads against their reference sequences manually to obtain an alignment and sequence identity rate for every read. For this we used Biopython[30] pairwise2.globalms with the same alignment parameters as minimap2 (matching score of 2, mismatch penalty of 4, gap open penalty of 4 and gap extension penalty of 2). The alignment string was parsed to compute *M*, *I*, *D* and *S* then accuracy and total error rate computed as above.

## 3 Results

### 3.1 Efficient Decoding of Variable-Length Nanopore Signals

A comparison of basecalling using global or chunk-level decoding is provided in **Table 1**. Basecalling using global decoding was significantly faster than chunk-level decoding, as expected if we examine the time complexity of each method. Observing the pseudocode in **Algorithm 1**, we see that given an input matrix *m* of size *S* × *T*, where *S* is the number of CTC symbols and *T* is the number of timesteps, the CTC beam search algorithm iterates *T* times. For each timestep, the *W* most probable beams are chosen and each extended by *S* symbols. Assuming the sorting algorithm has time complexity *O*(*N · log*(*N*)), the CTC beam search time complexity is *O*(*T · W · S · log*(*W · S*)). The global decoding approach also requires assembling the chunk-level CTC matrices, which has time complexity *O*(*T*), thus the global decoding time complexity is *O*(*T · W · S · log*(*W · S*) + *T*) = *O*(*T · W · S · log*(*W · S*)). The chunk-level decoding approach repeats the CTC beam search for every overlapping chunk and therefore considers each timestep up to *M* times, where *M* is the maximum number of chunks a timestep appears in, depending on the chunk length and step size chosen during segmentation. Thus the time complexity of local decoding is *O*(*T · W · S · log*(*W · S*) *· M*).

**Table 1:** Basecalling performance using chunk-level or global decoding and Biopython.pairwise2 alignment.

| Decoding approach | Used mRNA model? | Accuracy (Median) | Total error % (Median) | Median inference time per read (s) |
| --- | --- | --- | --- | --- |
| Chunk-level | No | 67.60 | 32.40 | 60.0 |
| Global | No | 67.90 | 32.10 | 17.3 |

Alignment of the basecalled sequences to the transcriptome with minimap2 resulted in differing numbers of mapped reads between the two decoding approaches, so accuracy was assessed using the Biopython.pairwise2 alignment to the reference sequence for every read. Global decoding achieved a small improvement in basecalling accuracy, consistent with our hypothesis that precise assembly of CTC matrices removes the ambiguity in determining the overlap of nucleotide sequences. Alignment results using minimap2 and Biopython.pairwise2 are detailed in **??**.

### 3.2 Sequence Biases in mRNA

The distribution of k-mer counts in the human transcriptome is provided in **Fig. 3**, along with some example 12-mer model estimations for three different sequence contexts. As expected, the distribution of k-mer counts in the transcriptome was nonuniform, suggesting a k-mer model holds some discriminatory power for estimating the most likely next nucleotide during basecalling.

**Figure 3:**
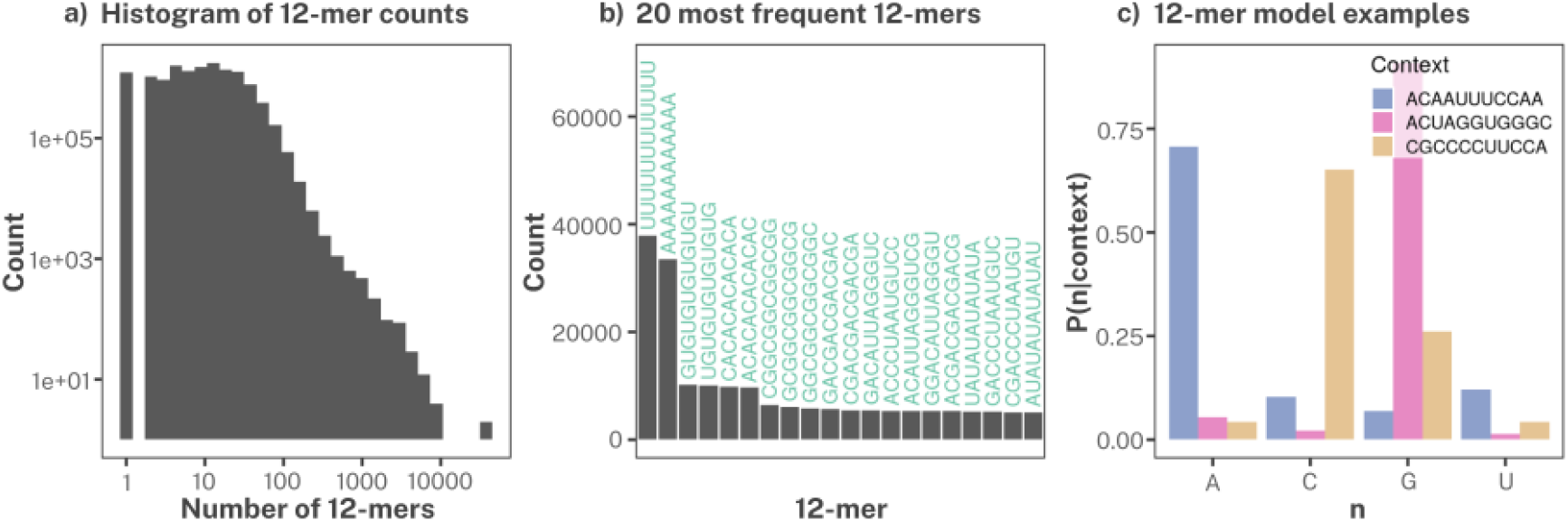
Visualisation of k-mer counts and mRNA model for *k* = 12. **a)** Histogram of 12-mer counts in the human transcriptome. **b)** The top 20 most frequent 12-mers (3’ to 5’) in the human transcriptome. **c)** mRNA model examples for three different sequence contexts.

### 3.3 Dynamic Inclusion of mRNA Language Model During CTC Decoding

Evaluation of the end-to-end basecaller on the independent GM12878 test set showed that inclusion of the mRNA model provides a small improvement in accuracy compared to basecalling without the mRNA model. Basecalling performance using the minimap2 alignment approach is summarized in **Table 2**. As can be seen in **Fig. 4**, when the CTC matrix output by the Sig2Seq model assigns similar probabilities to multiple nucleotides at the same timestep, the mRNA model can assist in selection of the correct nucleotide during decoding given the preceding sequence context.

**Table 2:** Basecalling performance with and without the mRNA model using minimap2 alignment.

| Decoding approach | Used mRNA model? | k-mer length | Signal entropy threshold | RNA model entropy threshold | Accuracy (Median) | Total error % (Median) |
| --- | --- | --- | --- | --- | --- | --- |
| Global | No | - | - | - | 77.42 | 22.58 |
| Global | Yes | 12 | 0.5 (medium) | 0.5 (medium) | 78.17 | 21.83 |

**Figure 4:**
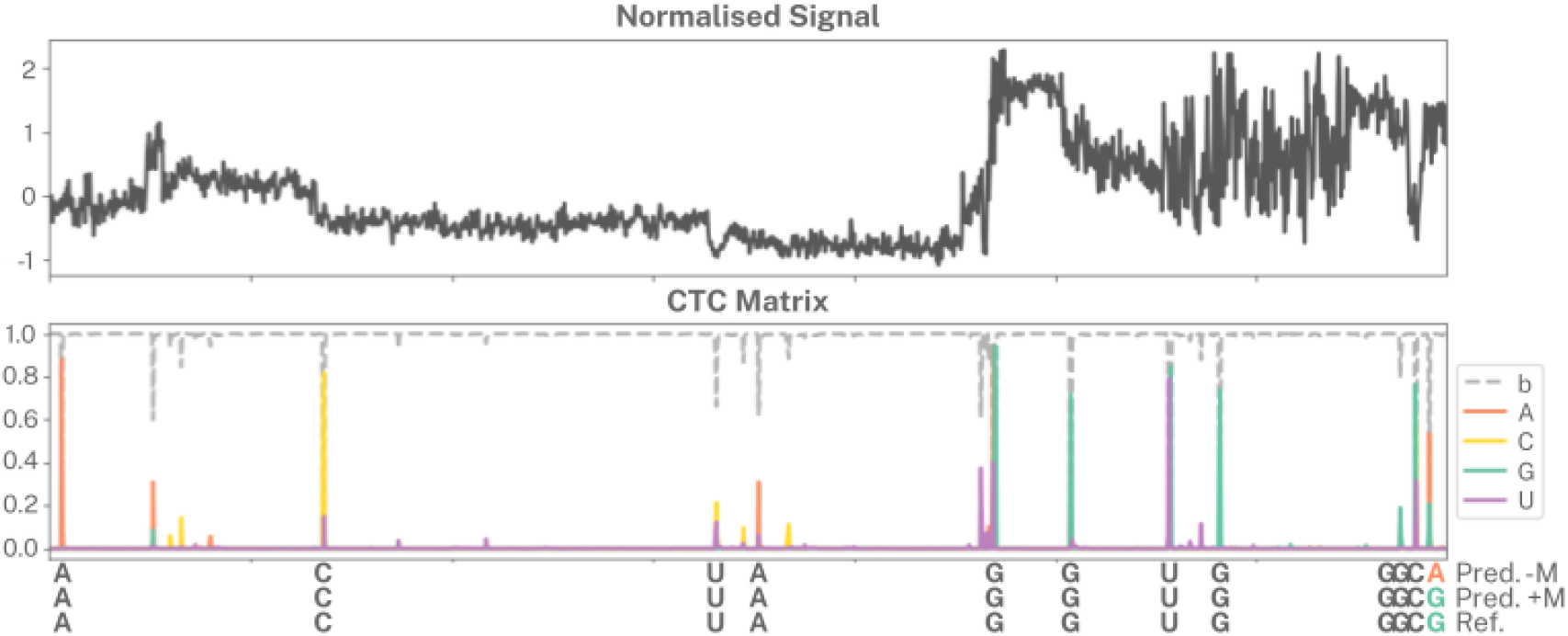
Visualization of the effect of the mRNA model on CTC beam search decoding. The normalized signal segment (upper panel) was fed to the Sig2Seq model, which generated the CTC matrix comprising a probability distribution across the symbols {*A, C, G, U, b*} for each timestep (visualised in the middle panel as a grid of probabilities and in the lower panel as a series of spikes). The nucleotide sequences obtained from decoding the CTC matrix without the mRNA model (Pred. −M) and with the mRNA model (Pred. +M), respectively, are listed below along with the reference sequence (Ref.). In this example, the mRNA model resolves the substitution error at the last nucleotide. The Sig2Seq model outputs similar probabilities for *A* and *G*, but the mRNA model favours inclusion of the correct nucleotide *G* given the context *ACUAGGUGGGC* (see **Fig. 3c**)

It is important to note that minimap2 failed to align a large number of reads in both tests, which we expect is because the error rate of the Sig2Seq model is beyond what minimap2 can tolerate. Since the focus of this study was our novel language-informed CTC decoding algorithm, the Sig2Seq model was not heavily optimized and so its error rate is unsurprising. It is therefore also pertinent to consider basecalling performance using the Biopython.pairwise2 alignment approach, which aligns every read to the reference sequence we computed based on the initial Guppy basecall alignment. The Biopython.pairwise2 alignment again showed that the mRNA model provides a small improvement to basecalling performance, supporting the minimap2 alignment results. The basecalling performance is detailed for both the minimap2 and Biopython.pairwise2 alignment approaches in **??**.

## 4 Discussion

In this study, we investigated whether inclusion of mRNA language information during CTC beam search decoding can be used to improve the accuracy of nanopore dRNA basecalling. We harnessed the sequence biases in mRNA language through a probabilistic k-mer model of the protein-coding transcriptome and proposed a novel approach to conditionally incorporate a language model during CTC beam search decoding. We showed that the mRNA language model is able to guide the beam search towards the more probable nucleotide when there is sufficiently high entropy in the probability distribution output by the Sig2Seq model, and overall results in a modest improvement to basecalling accuracy for mRNA.

We also proposed a novel way of combining chunk-level CTC matrices through averaging overlapping rows in each chunk to assemble a global matrix prior to CTC beam search decoding, rather than decoding each chunk individually and then assembling the resultant partial reads in nucleotide space. We showed both empirically and through analysis of the time complexity of each algorithm that compared to the chunk-level approach, global decoding is more efficient. Empirical results also suggest our approach reduces the error rate of the final nucleotide sequence, which we expect is because chunk-level assembly is exact in matrix space but ambiguous in nucleotide space. This approach can be applied to any basecalling algorithm for RNA or DNA that handles the varying length of nanopore signals through segmentation into chunks prior to inference. Importantly, our global decoding approach also facilitates the provision of long sequence contexts to a language model, which would be far less effective with a chunk-level approach since the language model could not be utilised in each chunk until a sufficiently long context had been decoded, or at all for nucleotide sequences shorter than the context.

It is important to acknowledge the potential limitations of modelling complex biological sequences with a simple k-mer model. Individual variability and pathological contexts can give rise to mRNA sequence variations that may not be represented in the reference transcriptome. Single nucleotide polymorphisms (SNPs) are estimated to occur at a rate of 5-8 per 10kB[31]. Although it is important to correctly call the SNP, given that the mRNA model is only included conditionally at timesteps where the signal model contains sufficient uncertainty, the likelihood of the mRNA model misdirecting the beam search towards the canonical nucleotide in favour of the variation is low but still possible. In the case of transcripts with altered or novel splicing patterns, since the context required by the proposed mRNA model is only 12 nucleotides it is likely to be fully contained within a single exon, which are on average 311 bp long[32]. Thus any splicing alteration other than intron retention is unlikely to affect the mRNA model context.

We think that these features give rise to an intriguing avenue of research beyond this study. Firstly, the mRNA model may better capture biological variability through the inclusion of SNPs and other known sequence variations in the training data, or even RNA modifications. Similarly, noise could be incorporated into the mRNA model training data to improve the robustness of the model to errors in the context sequence. In this case, neural language models may be better suited to capturing sequence variability beyond the training data and also facilitating longer context lengths to capture longer-range nucleotide dependencies. Finally, while the number of human protein-coding genes has converged to approximately 20,000, the corresponding set of protein-coding transcripts may not be complete [17]. Thus, as this information is uncovered, there is the opportunity to strengthen the mRNA model training data for enhanced discovery of new mRNAs. This study provides the first framework for language-informed dRNA basecalling, which lays the foundations for further dedicated RNA basecalling developments.

## Author Contributions

E.E. and A.S. conceived the study. E.E. supervised the project. A.S. designed and implemented the end-to-end basecalling architecture, including all data preparation, model development, testing and software implementation. A.S. devised the modified CTC beam search algorithm with input from E.E. N.S. performed the HEK293 sequencing. A.S. and E.E. contributed to the interpretation of results. A.S. wrote the manuscript with input from E.E and N.S.

