## Supplementary Material for "Language-Informed Basecalling Architecture for Nanopore Direct RNA Sequencing"

**Alexandra Sneddon**
Australian National University

**Nikolay Shirokikh**
Australian National University

**Eduardo Eyras**
Australian National University

#### Supplementary Material

#### S1 HEK293 Data Collection

##### S1.1 Cell Material

###### S1.1.1 Cell Lines and Culturing

HEK293T (human embryonic kidney) cells were purchased from American Type Culture Collection (ATCC). HEK293T cells were supplier-certified and grown in DMEM media supplemented with 10% FBS. Cells were cultured in humidity-controlled incubator at 37°C, 5% carbon dioxide ($CO_2$). The media was replaced every 72 hours, passaging and collection were done in T175 flasks at 80–90% confluency.

###### S1.1.2 Cell Pellet Collection

The HEK293 cells were collected from two T175 flasks when at 80-90% confluency using trypsin detachment. 5 ml of trypsin/EDTA were used to substitute the media, and flasks were incubated at 37°C for 5 minutes. The trypsinisation was ceased when cells were unadhered from the flasks by resuspending in 15 ml of complete DMEM media (with FBS). The suspension was then centrifuged at 200 g for 5 minutes at 4°C. The supernatant was discarded and cell pellets for each cell line were resuspended in 20 ml of phosphate buffered saline (PBS) with $Ca^{2+}$/$Mg^{2+}$. The suspensions were spun down as above. The supernatant was discarded and the PBS washing step was repeated. Residual PBS was removed, cell pellets were stored at -80°C and subsequently used for RNA extraction.

##### S1.2 RNA Extraction

###### S1.2.1 Cell Lysis

Cell lysis and polyadenylated RNA extraction were generally performed as described previously, with minor alterations [1]. Briefly, upon removal from -80°C, cell pellets were thawed on ice for 10 minutes prior to RNA extraction. Cell pellets were then lysed in 1 ml of denaturing lysis and binding buffer (100 mM Tris-HCl pH 7.4, 1% w/v lithium dodecyl sulfate (LiDS), 0.8 M lithium chloride, 40 mM EDTA and 8 mM DTT; LBB) by rigorous pipetting. For RNA extraction from total cell lysates, $5x10^{6}$ cells were lysed in 350 µl of RA1 lysis buffer (Macherey Nagel) and the RNA isolated according to manufacturer's instructions including an on-column DNase digestion step. RNA isolated from total cell lysate was then eluted from the columns in 80 µl of RNase-free water and stored at -80°C.

###### S1.2.2 Extraction of Polyadenylated RNA

500 µl of oligo(dT)25 magnetic beads (New England Biolabs) suspension was used per replicate for each cell pellet. The beads were washed with 1 ml of LBB twice, each time collecting the beads on a magnet and completely removing the supernatant. Upon washing, the oligo(dT)25 beads were resuspended in the pellet/LBB mixture and placed in a rotator set for 20 rpm at 25 °C for 5 minutes, followed by the same

rotation at 4 °C for 30 minutes. The suspension was briefly spun down at 12,000 g, separated on a magnet, and the supernatant was discarded. The beads were then resuspended in 1 ml wash buffer (20 mM Tris-HCl pH 7.4, 0.2% v/v Titron X-100, 0.4 M lithium chloride, 10 mM EDTA and 8 mM DTT; WB) and washed on a rotator set for 20 rpm at 4°C for 5 minutes, 3 wash rounds in total. The beads were collected on a magnetic rack and the supernatant was discarded. The wash procedure was repeated three times. The elution was carried out stepwise. Washed bead pellet was first resuspended in 50 µl of the elution buffer (25 mM HEPES-KOH, 0.1 mM EDTA; HE). The first suspension was heated at 60°C for 5 minutes to facilitate the elution, and the eluate was collected upon placing the bead-sample mixture on a magnetic rack, separating the beads, and recovering the clean supernatant. The resultant pellet was next resuspended in another 50 µl of HE buffer, and the process was repeated. The eluates were then combined and subjected to an additional solid-phase reversible immobilization (SPRI) bead purification step and stored frozen at -80°C.

###### S1.2.3 RNA Cleanup with SPRI Beads

The eluate from oligo(dT) bead extraction or in vitro polyadenylated RNA was further purified using AMPure XP SPRI beads (Beckman Coulter Life Sciences) according to the manufacturer's recommendations. Briefly, the eluate samples were supplemented with 1.2x volumes of the SPRI bead suspension in its standard (supplied) binding buffer, and the resultant mixture was incubated at room temperature for 5 minutes with periodic mixing. The SPRI beads were brought down by a brief 2,000 g spin down and separated from the solution on a magnetic rack. The supernatant was removed, and the beads were resuspended in 1 ml of 80% v/v ethanol, 20% v/v deionized water mixture and further washed by tube flipping. The bead and solution separation procedure were repeated. The ethanol washing process was repeated one more time. Any remaining liquid was brought down by a brief spin and removed using a pipette, and the beads were allowed to air-dry while in the magnetic rack for 2 minutes. The purified RNA was then eluted in 20 µl of deionized water and the RNA content was assessed using absorbance readout via Nanodrop and fluorescence-based detection via Qubit RNA high sensitivity (HS) assay kit (Thermo Fisher Scientific).

##### S1.3 MinION Sequencing

###### S1.3.1 dRNA Library Preparation and Loading

The flow cell priming and library sequencing protocol was followed as previously described (Simultaneous identification of m6A and m5C reveals coordinated RNA modification at single-molecule resolution | bioRxiv, no date)[1]. Briefly, 500-1,000 ng of SPRI-purified RNA from HeLa and HEK293T cells were used for each 2x library preparation within every replicate (all ONT-recommended volumes doubled) with direct RNA sequencing kit (SQK-RNA002) as supplied by ONT. SuperScript IV RNA Polymerase (Thermo Fisher Scientific) was used, RNA Control Standard (RCS) was omitted, and RNasin Plus (Promega) was included at 1 U/µl in all reaction solutions until the SPRI purification step after the reverse transcription reaction, as before. The final adaptor-ligated sample was eluted in 40 µl.

###### S1.3.2 dRNA Sequencing Run

Nanopore sequencing was conducted on an ONT MinION Mk1B using R9.4.1 flow cells for 72 hours. Initially, the flow cell was left at 25°C for 30 minutes to reach ambient temperature. The flow cell was inserted into the MinION Mk1B and a quality check was performed to ensure that the pore count was above manufacturer warranty level (800 pores). Prior to sample loading, the priming solution (Flush Buffer + Flush Tether) was degassed in a vacuum chamber for 5 minutes. A similar approach was used when loading the RNA library.

The run set up on the loaded libraries was performed using the default MinKNOW software (Version 4.5.25) run configuration. The SQK-RNA002 sequencing option was selected. For real-time assessment of the quality of the run, the output FAST5 files were basecalled inline with sequencing using Guppy in 'fast' basecalling mode.

#### S2 Sig2Seq Model Training Data Preparation

Fast5 files were basecalled with the ONT basecaller Guppy (v4.0.14) with options *–flowcell FLO-MIN106 –kit SQK-RNA002* and mapped to the reference transcriptome (GRCh38.p13 release 34) by minimap2[2] (v2.17) with options *-ax map-ont –secondary=no -t 15*. To retain only high confidence mappings, alignments that were either unmapped, mapped to the reverse strand or were secondary or supplementary mappings were removed using samtools[3] with option *-F 2324*. 10% of the remaining reads from each run were reserved for testing. The training reads were then "re-squiggled" using Nanopolish eventalign [4] with options *–samples –signal-index*, which aligns the signal to the reference sequence per 5-mer. To retain only signals with high confidence labels, reads containing more than 3 consecutive unaligned 5-mers were discarded, leaving 67,540 reads. The remaining signals were normalized using median absolute deviation with outlier smoothing, then segmented into overlapping chunks of length 1024 with step size 128 and labelled according to the Nanopolish alignment.

#### S3 Decoding Hyperparameter Optimization

##### S3.1 Approach

We tested several hyperparameter configurations for our modified CTC beam search decoder. We first compared our global decoding approach to the chunk-level decoding approach previously described, using 2,500 randomly selected reads in the reserved heart test set. For assembling the partial reads corresponding to each chunk, we used a simple pairwise assembly implementation from [5].

We also tested incorporating each of the mRNA models (6-mer, 12-mer) across three entropy thresholds and compared this to decoding without an mRNA model. We randomly selected 5,000 reads in the reserved heart test set that aligned to mRNA transcripts to use for testing. The entropy thresholds tested were:

**Table S1:** Tested entropy thresholds.

| Name | Signal entropy threshold | mRNA model entropy threshold |
|---|---|---|
| None | 0 | 1 |
| Medium | 0.5 | 0.5 |
| High | 0.9 | 0.1 |

Note that the entropy threshold 'None' means that the mRNA model is incorporated at every step. For all tests, a beam width of 6 was used. The test reads were basecalled end-to-end following the workflow depicted in **Fig. ??**. The sequence identity and total error rates were computed as in **??**.

#### S3.2 Global vs Chunk-Level Decoding Results

##### S3.2.1 Alignment With Minimap2

**Table S2:** Hyperparameter optimization results on 2,500 protein-coding reads from reserved heart test set.

| Decoding approach | Used mRNA model? | k-mer length | Signal entropy threshold | RNA model entropy threshold | # Mapped | Accuracy (Median) | % Insertions (Median) | % Deletions (Median) | % Substitutions (Median) | Total error % (Median) |
|---|---|---|---|---|---|---|---|---|---|---|
| Chunk-level | No | - | - | - | 117 | 82.61 | 3.28 | 9.24 | 4.59 | 17.39 |
| Global | No | - | - | - | 120 | 82.10 | 3.15 | 9.63 | 4.53 | 17.90 |

##### S3.2.2 Alignment With Biopython.pairwise2

**Table S3:** Hyperparameter optimization results on 2,500 protein-coding reads from reserved heart test set.

| Decoding approach | Used mRNA model? | k-mer length | Signal entropy threshold | RNA model entropy threshold | Accuracy (Median) | % Insertions (Median) | % Deletions (Median) | % Substitutions (Median) | Total error % (Median) |
|---|---|---|---|---|---|---|---|---|---|
| Chunk-level | No | - | - | - | 67.60 | 4.82 | 18.18 | 8.29 | 32.40 |
| Global | No | - | - | - | 67.90 | 4.80 | 17.85 | 8.25 | 32.10 |

#### S3.3 Modified CTC Beam Search Hyperparameter Optimization Results

##### S3.3.1 Alignment With Minimap2

**Table S4:** Hyperparameter optimization results on 5,000 protein-coding reads from reserved heart test set.

| Decoding approach | Used mRNA model? | k-mer length | Signal entropy threshold | RNA model entropy threshold | # Mapped | Accuracy (Median) | % Insertions (Median) | % Deletions (Median) | % Substitutions (Median) | Total error % (Median) |
|---|---|---|---|---|---|---|---|---|---|---|
| Global | No | - | - | - | 233 | 82.96 | 3.21 | 9.60 | 4.30 | 17.04 |
| Global | Yes | 6 | 0 (none) | 1 (none) | 239 | 83.02 | 3.20 | 9.52 | 4.30 | 16.98 |
| Global | Yes | 6 | 0.5 (medium) | 0.5 (medium) | 239 | 83.02 | 3.20 | 9.52 | 4.30 | 16.98 |
| Global | Yes | 6 | 0.9 (high) | 0.1 (high) | 239 | 83.02 | 3.20 | 9.52 | 4.30 | 16.98 |
| Global | Yes | 12 | 0 (none) | 1 (none) | 205 | 81.99 | 3.83 | 9.14 | 5.16 | 18.01 |
| Global | Yes | 12 | 0.5 (medium) | 0.5 (medium) | 239 | 83.03 | 3.21 | 9.35 | 4.31 | 16.97 |
| Global | Yes | 12 | 0.9 (high) | 0.1 (high) | 240 | 82.94 | 3.21 | 9.53 | 4.32 | 17.06 |

##### S3.3.2 Alignment With Biopython.pairwise2

**Table S5:** Hyperparameter optimization results on 5,000 protein-coding reads from reserved heart test set.

| Decoding approach | Used mRNA model? | k-mer length | Signal entropy threshold | RNA model entropy threshold | Accuracy (Median) | % Insertions (Median) | % Deletions (Median) | % Substitutions (Median) | Total error % (Median) |
|---|---|---|---|---|---|---|---|---|---|
| Global | No | - | - | - | 67.50 | 5.39 | 17.55 | 8.20 | 32.50 |
| Global | Yes | 6 | 0 (none) | 1 (none) | 67.49 | 5.39 | 17.55 | 8.20 | 32.51 |
| Global | Yes | 6 | 0.5 (med.) | 0.5 (med.) | 67.48 | 5.38 | 17.54 | 8.21 | 32.52 |
| Global | Yes | 6 | 0.9 (high) | 0.1 (high) | 67.50 | 5.38 | 17.55 | 8.21 | 32.50 |
| Global | Yes | 12 | 0 (none) | 1 (none) | 67.37 | 5.73 | 17.17 | 8.45 | 32.63 |
| Global | Yes | 12 | 0.5 (med.) | 0.5 (med.) | 67.57 | 5.40 | 17.53 | 8.21 | 32.43 |
| Global | Yes | 12 | 0.9 (high) | 0.1 (high) | 67.49 | 5.38 | 17.55 | 8.20 | 32.51 |

### S4 Decoding on Independent Test Set

#### S4.1 Alignment With Minimap2

**Table S6:** Decoding results on independent test set with and without mRNA model.

| Decoding approach | Used mRNA model? | k-mer length | Signal entropy threshold | RNA model entropy threshold | # Mapped | Accuracy (Median) | % Insertions (Median) | % Deletions (Median) | % Substitutions (Median) | Total error % (Median) |
|---|---|---|---|---|---|---|---|---|---|---|
| Global | No | - | - | - | 73 | 77.42 | 2.70 | 13.63 | 6.15 | 22.58 |
| Global | Yes | 12 | 0.5 (medium) | 0.5 (medium) | 73 | 78.17 | 2.70 | 13.84 | 5.83 | 21.83 |

#### S4.2 Alignment With Biopython.Pairwise2

**Table S7:** Decoding results on independent test set with and without mRNA model.

| Decoding approach | Used mRNA model? | k-mer length | Signal entropy threshold | RNA model entropy threshold | Accuracy (Median) | % Insertions (Median) | % Deletions (Median) | % Substitutions (Median) | Total error % (Median) | |
|---|---|---|---|---|---|---|---|---|---|---|
| Global | No | - | - | - | 73 | 66.12 | 4.11 | 20.34 | 8.51 | 33.88 |
| Global | Yes | 12 | 0.5 (medium) | 0.5 (medium) | 73 | 66.15 | 4.13 | 20.27 | 8.52 | 33.85 |
